# Impacts of Perinatal Nicotine Exposure on nAChR Expression and Glutamatergic Synaptic Transmission in the Mouse Auditory Brainstem

**DOI:** 10.1101/2024.05.08.592930

**Authors:** Mackenna Wollet, Abram Hernandez, Kaila Nip, Jason Pugh, Jun Hee Kim

## Abstract

Exposure to nicotine in utero, often due to maternal smoking, significantly elevates the risk of auditory processing deficits in offspring. This study investigated the effects of chronic nicotine exposure during a critical developmental period on the functional expression of nicotinic acetylcholine receptors (nAChRs), glutamatergic synaptic transmission, and auditory processing in the mouse auditory brainstem. We evaluated the functionality of nAChRs at a central synapse and explored the impact of perinatal nicotine exposure (PNE) on synaptic currents and auditory brainstem responses (ABR) in mice. Our findings revealed developmentally regulated changes in nAChR expression in the medial nucleus of the trapezoid body (MNTB) neurons and presynaptic Calyx of Held terminals. PNE was associated with enhanced acetylcholine-evoked postsynaptic currents and compromised glutamatergic neurotransmission, highlighting the critical role of nAChR activity in the early stages of auditory synaptic development. Additionally, PNE resulted in elevated ABR thresholds and diminished peak amplitudes, suggesting significant impairment in central auditory processing without cochlear dysfunction. This study provides novel insights into the synaptic disturbances that contribute to auditory deficits resulting from chronic prenatal nicotine exposure, underlining potential targets for therapeutic intervention.

## Introduction

Prenatal nicotine exposure, resulting from maternal smoking during pregnancy, has been shown to cause a range of long-term neurodevelopmental defects in children^1–5^, including attention-deficit disorder and learning disabilities which are shown to be comorbid with sensory processing deficits^3–5^. Due to the increased prevalence of electronic cigarettes, this begs the need for investigation into the effects of prenatal exposure to nicotine alone on fetal health and development^6^. In utero nicotine exposure can cause hearing impairments and auditory processing disorders leading to speech and language delays, poor sound localization, and difficulty understanding speech in noisy environments^1^. Animal studies have demonstrated that nicotine exposure during development impairs temporal processing in an auditory startle test and disrupts development of glutamatergic synapses in the auditory cortex^7,8^. Nicotine is known to bind to and activate nAChRs, which are widely distributed throughout the auditory nervous system and play a key role in synaptic transmission^9–11^. However, the specific effects of developmental nicotine exposure on nAChR signaling and auditory processing are not yet fully understood. In this study, we aim to investigate the impact of nicotine exposure during early development on auditory brainstem nAChR signaling, synaptic transmission, and auditory processing.

nAChRs are expressed throughout the auditory system and play a crucial role in auditory processing. These receptors are found on several peripheral and central auditory cells. Within the cochlea, α*9* nAChRs are expressed on postsynaptic outer hair cells receiving cholinergic medial olivocochlear efferent terminals^12^. In the auditory brainstem, nAChR expression has been demonstrated with receptor autoradiography, in situ hybridization, and immunohistochemistry of rodent cochlear nucleus, superior olivary complex, and inferior colliculus^13–15^. Additionally, recent studies have confirmed the presence of nAChRs across all layers of the auditory cortex^11,16^, where cholinergic modulation of neuronal activity enhances responses to auditory inputs^17,18^. In the inferior colliculus, α3β4 nAChRs, a rarer nAChR subtype, play a role in regulating the excitability of VIP neurons^19^. Moreover, in vivo recordings of the MNTB showed that nAChRs play a critical role in regulating tone-evoked activity and accurately coding signal-in-noise detection^9^. In addition to nAChR expression, muscarinic acetylcholine receptors modulate excitability of MNTB principal neurons before hearing onset^20^.

In terms of subcellular location at central synapses, nAChRs can be located on both pre- and postsynaptic domains to regulate neurotransmission. For example, presynaptic α7 nAChRs increase glutamate release to enhance NMDAR-mediated excitatory postsynaptic potentials (EPSPs) in apical dendrites of pyramidal neurons in the developing auditory cortex, and this effect is eliminated by p21^11^. This transient effect highlights the importance of nAChRs during the critical period of development in the auditory cortex suggesting a role in glutamatergic synapse maturation. Postsynaptic nAChR expression in neurons of the ventral nucleus of the lateral lemniscus (VNLL) is observed before hearing onset and subsequently declines with age^21^. Despite the importance of nAChR expression at local synapses during auditory development, the cellular mechanisms by which chronic nicotine exposure during early postnatal development alters synaptic transmission at the level of a single synapse remain unclear.

The Calyx of Held-MNTB synapse is a very reliable synapse with high temporal precision that allows for binaural sound localization computations in the auditory brainstem. Studies utilizing autoradiography techniques have demonstrated the dynamic expression pattern of nAChRs in the MNTB, which is high during the second postnatal week, typically considered the critical period of auditory development^15^. Concurrently, PNE impairs development of glutamatergic inputs and temporal precision of action potential firing in the VNLL within the auditory brainstem^21^.

In this study, we employed a PNE model that subjects neonatal pups to nicotine (postnatal days 8-12) to investigate effects of developmental nicotine exposure during the critical period similar to the fetal in utero condition. Utilizing ex vivo and in vivo electrophysiology in this model, we aim to study the impact of PNE on the developmental processes of auditory synapses within the auditory brainstem. Specifically, we focus on investigating the functional expression of nAChRs within the Calyx of Held-MNTB synapse during the postnatal period and delineating how PNE perturbs the developmental dynamics of nAChRs and the maturation of glutamatergic transmission. By elucidating the cellular mechanisms whereby PNE impact auditory brainstem development and processing, this research contributes to uncover potential therapeutic targets and interventions aimed at ameliorating the adverse effects of PNE on auditory function.

## Results

### Functional expression of nAChRs at the Calyx-MNTB synapse during postnatal development

We investigated the expression of nAChRs and their activity at the Calyx-MNTB synapse during postnatal development. Previous work has demonstrated expression of α7 nAChRs within the MNTB peaks during the second postnatal week via in situ hybridization^15^. To assess the functional expression of nAChRs, we conducted whole-cell patch clamp recordings to measure ACh-evoked currents and membrane potential changes in MNTB principal neurons across postnatal development (from postnatal day 8, p8 to p20). In the presence of atropine (1 μM) to block muscarinic receptor effects, local application of acetylcholine (ACh, 1 mM) with short pressure puff (1 s) on the MNTB neuron induced ACh-evoked currents in voltage-clamp mode (holding at −65 mV). The average amplitude of ACh-evoked currents was −32.8 ± 7.48 pA (n=6) at p8, which significantly reduced to an average amplitude of −2.8 ± 10.99 pA (n=6) by p20 (**Figure 1A-B**; One-way ANOVA, p=0.0002, n=6-17 cells per age timepoint). Plotting of the amplitude against postnatal age was fit by linear regression illustrating a significantly non-zero slope (Slope=1.8, R^2^=0.48, **Supplemental Figure 1**; T-test, F=7.386, p=0.02). Our results revealed a significant decline in ACh-evoked inward currents in MNTB principal neurons during postnatal development.

**Figure 1.**
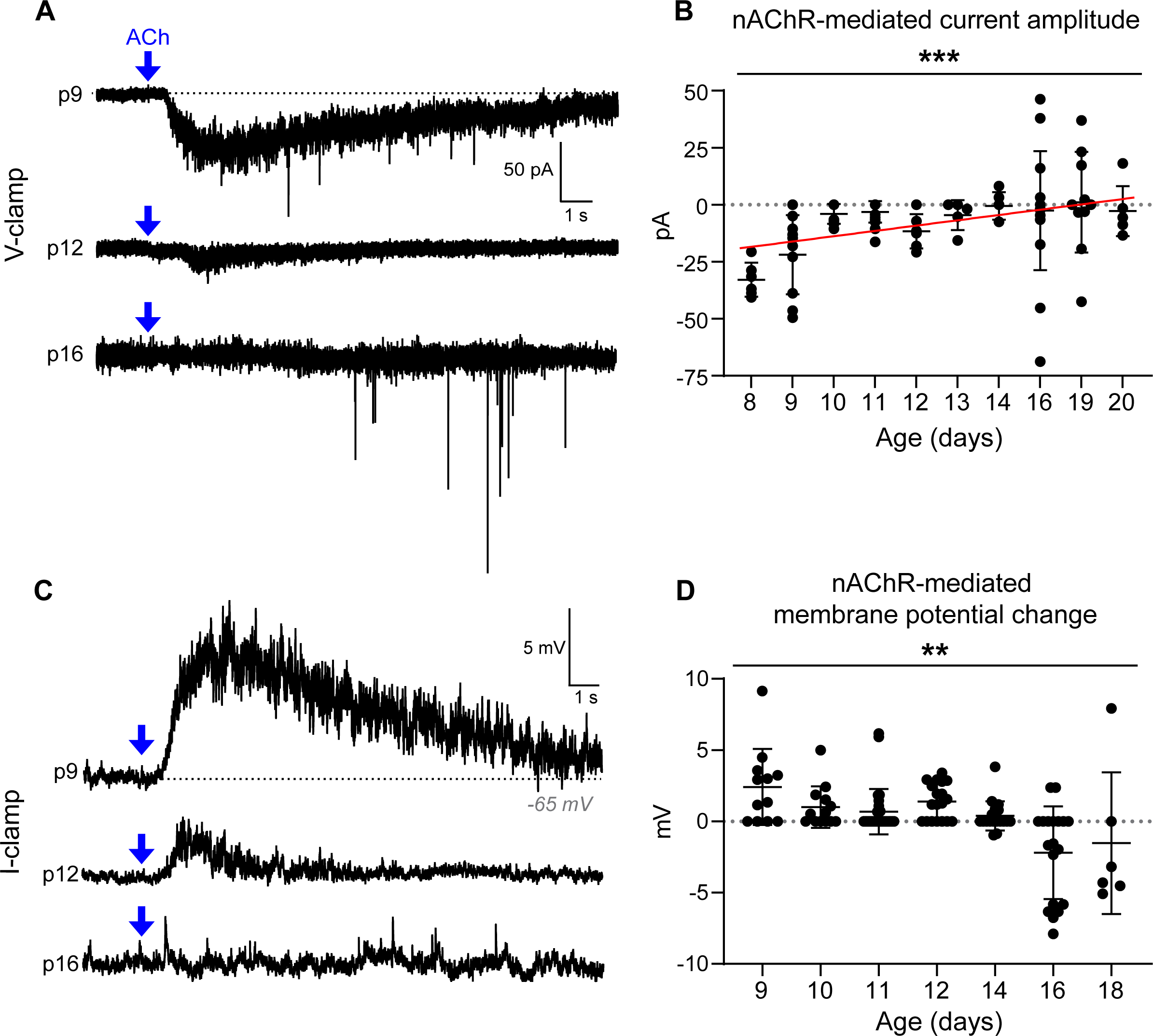
Age-dependent decrease in nAChR functional expression in MNTB principal neurons during postnatal development. **A.** Representative current traces illustrating nAChR-mediated responses in MNTB neurons at postnatal day 9 (p9), p12, and p16, recorded using voltage-clamp configuration (holding potential at −65 mV). ACh puffs (1 mM, 1 s) were administered at the timepoints denoted by blue arrows. **B.** Scatter plot displaying the amplitude of nAChR-mediated currents as a function of postnatal age up to p20. Each data point represents an individual neuron. A significant age-related decline in the amplitude of nAChR-mediated currents was observed (p=0.0002, One-way ANOVA). **C.** Representative traces depicting changes in the membrane potential of MNTB neurons in response to ACh puffs at p9, p12, and p16. **D.** Quantitative analysis of membrane potential changes, determined by calculating the difference between resting potential and peak potential following acute ACh application. ACh-evoked depolarization was found to be significantly reduced up to p18 (p=0.0029, One-way ANOVA). All quantitative data are presented as mean ± S.E.M.

In addition to measuring ACh-evoked currents, we also evaluated age-dependent changes in membrane potential in response to ACh puff. At p9, ACh puff application caused a consistent and clear depolarization of the membrane potential in MNTB neurons by +2.4 ± 2.66 mV (n=12). However, this depolarization was not sufficient to evoke action potentials in MNTB neurons. Like ACh-evoked currents, this ACh-evoked depolarization diminished over development and eventually resulted in hyperpolarization by p18 (−1.53 ± 4.97 mV, n=6). The statistical analysis revealed a significant change in ACh-evoked membrane potential change across postnatal development (**Figure 1C-D**; One-way ANOVA, p=0.003, 6-29 cells per age timepoint). A linear regression analysis revealed a significant non-zero slope when fitting membrane potential changes against postnatal age, indicating an age-dependent decline in functional nAChRs (**Supplemental Figure 1**, T-test, F=21.8, p=0.005). Taken together, the results highlight the developmental changes in functional nAChR expression in postsynaptic MNTB neurons.

Previous work has demonstrated the involvement of α*4*β*2* and α*7* nAChRs in the activity of MNTB neurons *in vivo*^9^. To determine the specific nAChR subtypes mediating ACh-evoked currents in MNTB neurons, we employed selective antagonists, DHβE and MLA, for α*4*β*2* and α*7* nAChRs, respectively. We recorded ACh-evoked currents in MNTB neurons from p9-10 mice to maximize current amplitudes. In control, repeated application of ACh (1 mM) at 9-minute intervals led to a decrease in the amplitude of ACh-evoked currents due to receptor desensitization (**Figure 2A, D**; Paired t-test: −15.65 ± 4.93 pA to −11.85 ± 4.25 pA, n=11 cells, p=0.0043). Desensitization of nAChRs in response to repeated agonist application is well documented^22,23^; thus, even in the absence of nAChR antagonists, ACh-evoked currents decrease in amplitude with subsequent ACh applications. To account for desensitization, we calculated the “paired-puff ratio”, (Current amplitude from second puff) / (Current amplitude from first puff) and compared it between drug treatments and control conditions. A significantly smaller paired-puff ratio in the presence of an antagonist indicates the involvement of the antagonized nAChR subtype in mediating ACh-evoked currents. Our results revealed an average paired-puff ratio of 0.71 ± 0.07 in the control aCSF condition. There was a significant effect of drug treatment on the paired-puff ratio across all treatments (One-way ANOVA, p=0.017, F=4.78). The paired-puff ratio is not significantly different between control and MLA treatments (**Figure 2E**; Dunnett’s multiple comparisons test, MLA: 0.65 ± 0.12, p=0.88, n=11 and 9). However, the paired-puff ratio is significantly smaller than control in DHβE condition (Control: 0.71 ± 0.07, DHβE: 0.26 ± 0.12, p=0.015, n=11 and 7). These data suggest that DHβE-sensitive receptors (α*4*β*2* nAChRs) contribute significantly to the postsynaptic currents observed in pre-hearing MNTB neurons.

**Figure 2.**
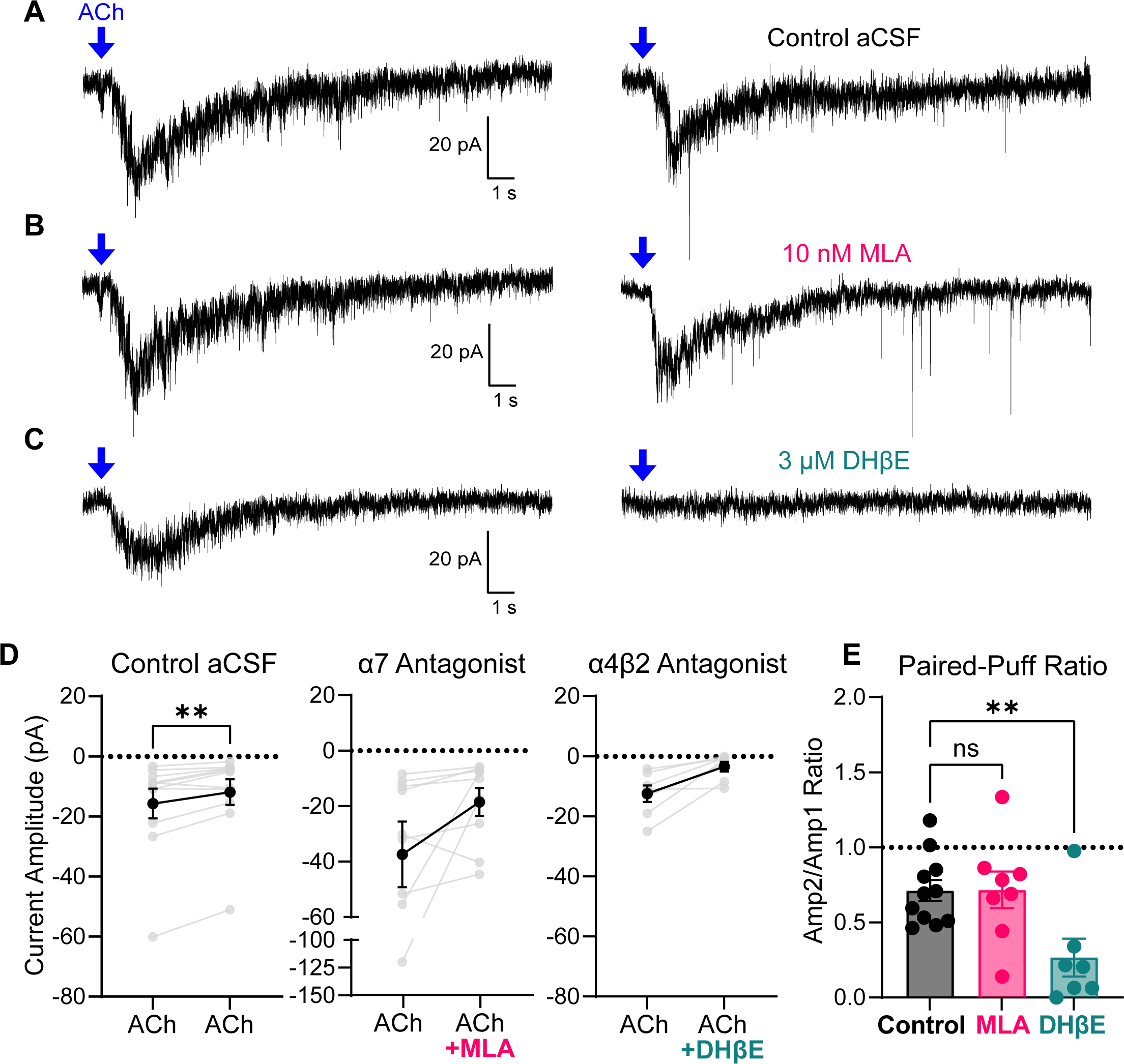
α4β2 nAChRs, not α7 nAChRs, mediate postsynaptic nAChR currents in pre-hearing MNTB principal neurons. **A-C.** Representative traces of postsynaptic ACh-evoked currents at p9-10 in the presence of atropine. ACh puff denoted by a blue arrow. Traces on right are second ACh puff applications with control aCSF or nAChR antagonist bath applied. **D.** Quantification of ACh-evoked currents pre- and post-bath application of drug. Individual data points are shown in gray and mean ± SEM is shown in black. In control aCSF, the first ACh-evoked current amplitude was significantly larger than the second (Paired t-test, n=11 cells, p=0.0043). **E.** The paired-puff ratio was calculated as follows: 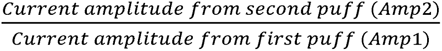. There was a significant effect of drug treatment on paired-puff ratio (p=0.017, F=4.78). Using Dunnett’s multiple comparisons test, DHβE had a significantly smaller paired-puff ratio compared to control (p=0.015) while MLA was not significantly different from control (p=0.89).

### The activation of nAChR enhances vesicular glutamate release from the calyx terminal

Upon postsynaptic recording from mice after p14, when postsynaptic ACh-evoked currents were almost diminished, we found an increase in spontaneous excitatory postsynaptic currents (sEPSCs) following ACh application. To investigate the presence of nAChR at presynaptic terminals and impact of its activation on presynaptic vesicular glutamate release, we recorded miniature EPSCs (mEPSCs) in MNTB neurons from post-hearing mice (p14 to p22). To avoid the effect of ACh-evoked muscarinic receptor activation at presynaptic terminal, we used atropine (1 μM) for all mEPSC experiments. In the presence of TTX (0.5 μM) and atropine (1 μM), we recorded mEPSCs from MNTB principal neurons before and after ACh puff. Compared to baseline, the frequency and amplitude of mEPSCs significantly increased following ACh application (**Figure 3A-B**; Wilcoxon matched-pairs test of Frequency: 0.8177 ± 0.27 to 3.455 ± 0.67 Hz, p<0.0001, n=32 cells; Amplitude: 43.22 ± 4.16 to 52.85 ± 4.98 pA, p=0.0069, n=33 cells). This indicates that presynaptic nAChR activation increases spontaneous vesicular glutamate release from the calyx terminal. Furthermore, the quantitative analysis of mEPSC amplitude using a cumulative frequency graph and a frequency distribution histogram, showing the rightward shift of mEPSC amplitude distribution following ACh application, support the increase of glutamatergic transmission at the calyx of Held synapse (**Figure 3C**). To examine if the effect of nAChR activation on glutamate release was dependent on calcium influx into presynaptic terminal, we performed the same experiments in a calcium-free extracellular condition using aCSF where CaCl_2_ was replaced with MgCl_2_ (0 mM CaCl_2_ and 3 mM MgCl_2_). In the absence of extracellular calcium, nAChR activation by ACh puff had no effect on mEPSC frequency and amplitude (**Figure 3D-E**; Frequency: p=0.16, n=11 cells; Amplitude: p=0.69 using Wilcoxon test, n=10 cells), indicating calcium flux is important for nAChR activation-mediated glutamate release at the Calyx terminal. We examined if ACh-evoked effects on mEPSCs are mediated by α*7* nAChRs, which has a high calcium permeability relative to other nAChR subtypes. We measured mEPSC frequency and amplitude in response to ACh in the presence of MLA, a selective α7 nAChR antagonist. With α*7* nAChRs antagonized, ACh puff had no effect on mEPSC frequency or amplitude (**Figure 3F-G**; Frequency: p=0.71, n=13 cells; Amplitude: p=0.79 n=12 cells using paired t-tests). The results suggest that after hearing onset, presynaptic α*7* nAChR activation enhances vesicular glutamate release at the Calyx terminal that is dependent on calcium flux into presynaptic terminal.

**Figure 3.**
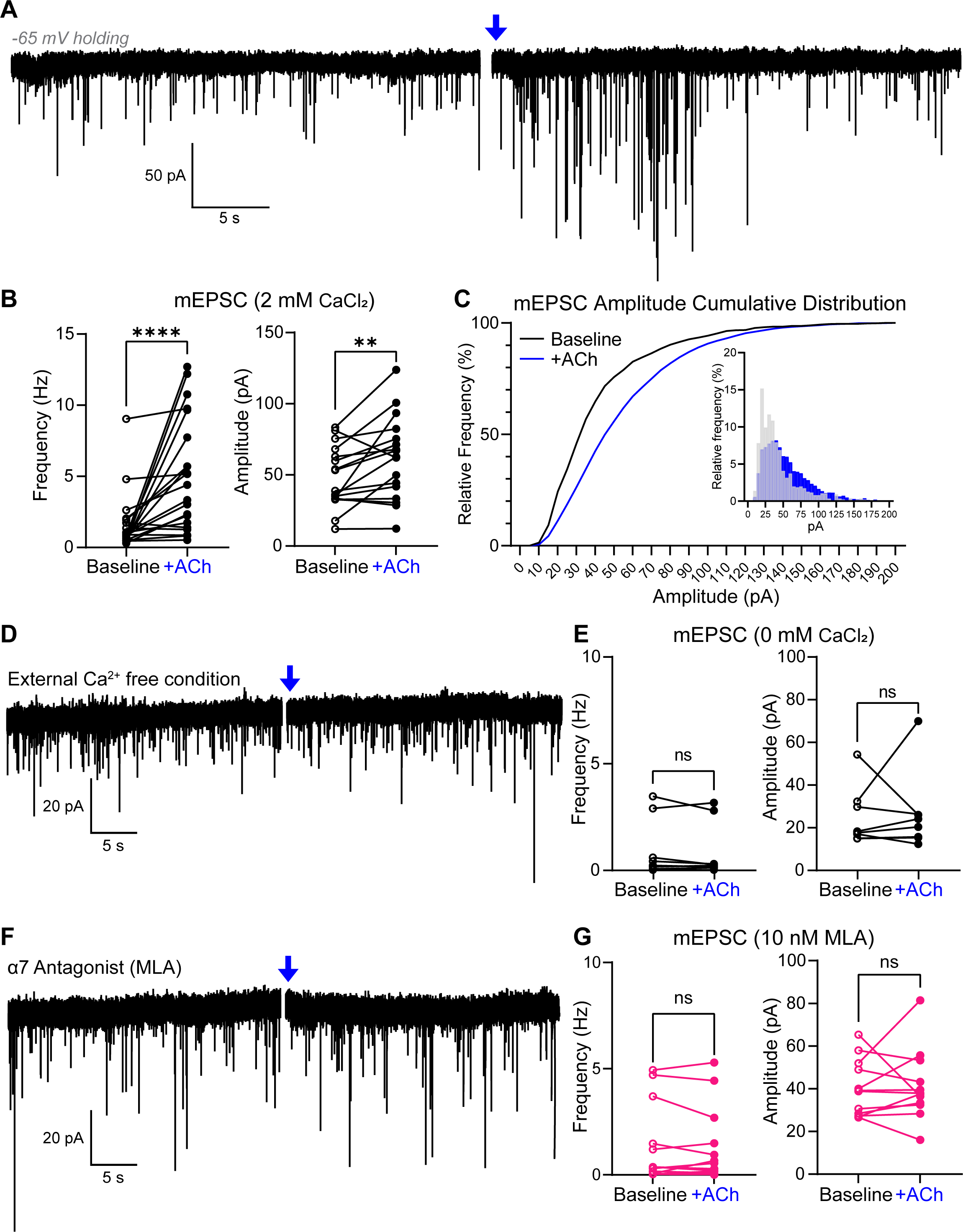
Activation of nAChR enhances vesicular glutamate release from the calyx of Held terminal. **A.** Representative trace of mEPSCs in an MNTB neuron (at p16) in the presence of TTX (0.5 μM). ACh puff application denoted by a blue arrow. **B.** mEPSC frequency and amplitude were quantified 30 seconds before (Baseline, black) and immediately after ACh application (+ACh, blue). Following ACh application, there was a significant increase in frequency (Wilcoxon matched-pairs rank test, p<0.0001, n=32 cells) and amplitude (Paired t-test, p=0.007, n=33 cells). **C.** Cumulative frequency distribution of mEPSC amplitudes at baseline and following ACh application. Inset: Frequency histogram displaying mEPSC amplitudes. **D.** Representative traces of mEPSCs in calcium-free external solution. **E.** Quantification of mEPSC frequency and amplitude before and after ACh application in calcium-free external solution. There was no significant difference between frequency (p=0.16, n=11 cells) or amplitude (p=0.69, n=10 cells) using Wilcoxon test. **F.** Representative traces of mEPSCs in presence of MLA, an α*7* nAChR antagonist. **G.** Quantification of mEPSC frequency and amplitude before and after ACh application in presence of MLA. No significant difference was observed post-ACh puff in frequency (p=0.71, n=13 cells) or amplitude (p=0.79, n=12 cells) using paired t-tests.

### PNE increases nAChR-mediated currents in postsynaptic MNTB neurons after hearing onset

Chronic nicotine exposure has been known to upregulate nAChR expression across brain regions^24–26^. To investigate the impact of chronic nicotine exposure on nAChR functional expression at pre- and postsynaptic terminal and synaptic function in the MNTB during the early stages of postnatal development, we established a perinatal nicotine exposure mouse model. In our PNE protocol, nicotine (0.7 mg/kg free base) or saline vehicle was administered to mouse pups (p8-p12, before hearing onset) twice daily via subcutaneous injections (**Figure 4A**). We evaluated postsynaptic nAChR-mediated currents during postnatal development between PNE and vehicle mice. Notably, PNE mice exhibited larger postsynaptic ACh-evoked currents after hearing onset compared to control. This effect was most pronounced at p20-p21 (**Figure 4B-C**, Two-way ANOVA: p=0.038, n=7-35 cells per timepoint; Sidak’s multiple comparisons between control and PNE at p20-21: p=0.001, n=15-22 cells). Chronic nicotine exposure before hearing onset (p8-p12) induced a sustained increase in ACh-evoked currents in MNTB neurons that persists through post-hearing ages. This result suggests that chronic nicotine exposure has a significant impact on the developmental patterning of nAChR expression in postsynaptic MNTB neurons in the auditory brainstem.

**Figure 4.**
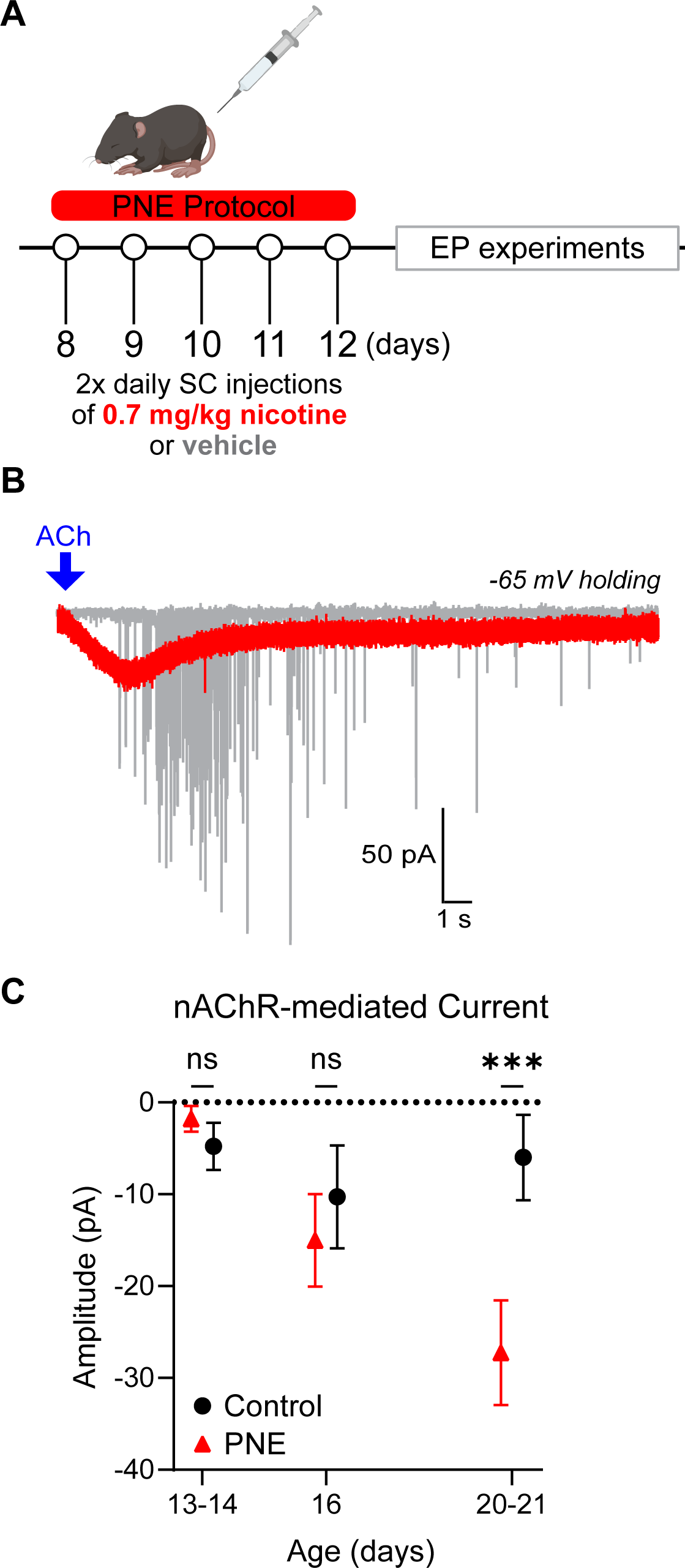
PNE augments postsynaptic nAChR-mediated currents in MNTB neurons following hearing onset. **A.** Schematic of nicotine injection protocol used to establish PNE model. **B.** Representative trace of nAChR-mediated currents following ACh application in control (gray) and PNE (red) neurons at p20. **C.** Quantification of ACh-evoked currents in postsynaptic MNTB neurons across ages in control and PNE animals. There is a significant effect of treatment on current amplitudes, and this effect is most pronounced at the p20-21 timepoint in PNE mice (Two-way ANOVA, p=0.038, n=7-35 cells per time point per group; Sidak’s multiple comparisons at p20-21: p=0.001, n=15-22 cells).

### PNE modifies presynaptic nAChR expression and its influence on glutamate release at the calyx terminal

In WT mice, nAChR activation by ACh puff enhances the spontaneous release of vesicular glutamate at the presynaptic terminal across post-hearing ages (**Figure 3**). Next, we investigated whether PNE impacts nAChR-mediated presynaptic glutamate release. mEPSC frequency and amplitude were analyzed in control and PNE mice (at p20) before (baseline) and after ACh application. At baseline, there was no significant difference in the frequency of mEPSCs between PNE and control mice (Control: 0.55 ± 0.13 Hz vs PNE: 0.70 ± 0.15 Hz, Mann-Whitney test, p=0.81, n=17-28 cells/group). However, the amplitude of mEPSC was significantly decreased in PNE mice compared to control (Control: 40.28 ± 3.07 pA vs PNE: 31.92 ± 2.35 pA, Welch’s t-test, p=0.039, n=15-29 cells/group), suggesting an impairment in the synaptic transmission in PNE mice (**Figure 5A-B**). In response to an ACh puff, control mice displayed a significant increase in both mEPSC frequency and amplitude (**Figure 5C**, Fisher LSD test, p<0.0001 and p=0.042, n=17 and 18 cells, respectively), whereas PNE mice showed no significant change in either mEPSC frequency (Fisher LSD test, p=0.67, n=28 cells) or amplitude (Fisher LSD test, p=0.88, n=28 cells) following acute ACh application. The results indicate that PNE disrupts the functional role of nAChR in regulation of neurotransmitter release at the presynaptic terminal. Furthermore, chronic nicotine exposure led to a reduction in baseline mEPSC amplitude in PNE mice, indicating detrimental effects of PNE on synapse development. Together, these findings suggest that PNE diminishes presynaptic nAChR expression and impairs synaptic transmission at the calyx-MNTB synapse.

**Figure 5.**
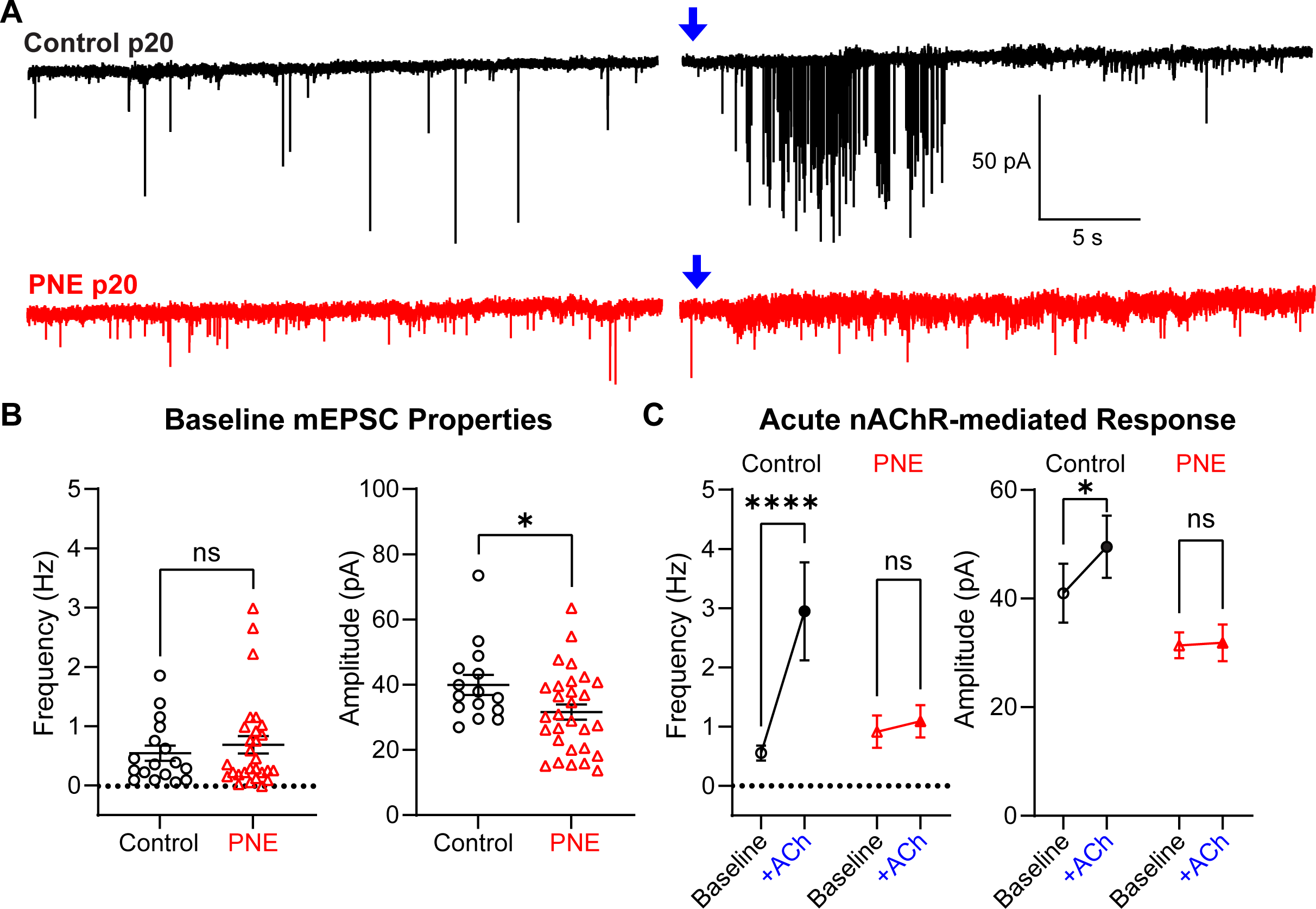
PNE alters glutamate release and nAChR-mediated enhancement in the calyx terminal. **A.** Representative traces of mEPSCs from control (black) and PNE mice (red) at p20. Blue arrows indicate the time of acute ACh application (1 mM, 1 s). **B.** Quantitative analysis of baseline mEPSC properties from the 30 seconds preceding ACh puff. There is no significant change in baseline mEPSC frequency between control and PNE mice (Mann-Whitney test, p=0.81, n=17-28 per group). However, a significant decrease in average baseline mEPSC amplitude is evident in PNE mice (Welch’s t-test, p=0.039, n=15-29 per group). **C.** Acute effects on mEPSCs following ACh application in control (black) and PNE mice (red). Like WT in Figure 2, vehicle control mice had increased mEPSC frequency and amplitude in response to ACh (Fisher LSD test, p<0.0001 and p=0.042, respectively, n=17-18 cells). In PNE mice, ACh puff had no significant impact on mEPSC frequency (Fisher LSD test, p=0.67, n=28 cells) or mEPSC amplitude (Fisher LSD test, p=0.88, n=28 cells).

### PNE impairs glutamatergic neurotransmission at the calyx-MNTB synapse

PNE alters baseline mEPSC properties in the absence of acute ACh application (Figure 5). To investigate the impact of PNE on glutamatergic neurotransmission at the calyx-MNTB synapse, we recorded evoked EPSCs in MNTB principal neurons using afferent fiber stimulation. We used strychnine (2 μM, glycine receptor antagonist) and bicuculline (20 μM, GABA_A_ receptor antagonist) to avoid the interference of inhibitory synaptic inputs. We found that PNE resulted in a significant decrease in the amplitude of single EPSCs recorded from MNTB principal neurons at p20. The amplitude of eEPSC in PNE mice was significantly decreased compared to control (**Figure 6A-B**, Control: 3.08 ± 0.30 nA vs. PNE: 1.54 ± 0.26 nA; Mann-Whitney, p=0.001, n=14 cells/group). To further evaluate release probability, we utilized a paired pulse protocol with a 50 ms interstimulus interval and quantified the paired-pulse ratio (PPR), which is the amplitude ratio of the second EPSC to the first EPSC (EPSC2/EPSC1). In PNE mice, PPR is significantly increased compared to control (**Figure 6C**, 0.96 ± 0.03 and 0.81 ± 0.02, respectively, Mann Whitney, p=0.0002, n=14 cells/group). These results suggest that PNE has a significant impact on presynaptic release probability and consequently synaptic transmission at the calyx-MNTB synapse.

**Figure 6.**
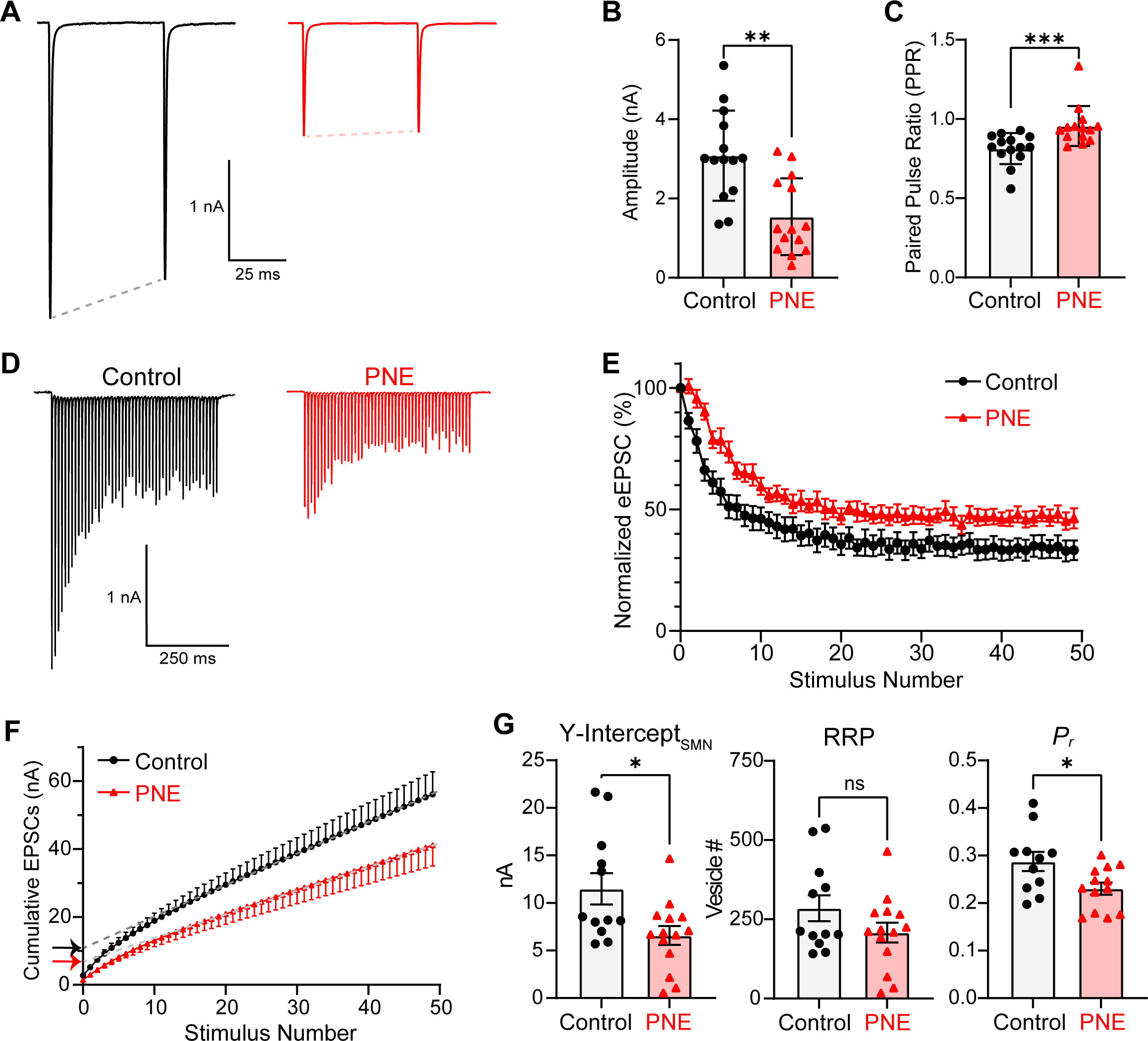
PNE impairs glutamatergic neurotransmission at the Calyx-MNTB synapse. **A.** Representative traces of evoked paired EPSCs (50 ms interstimulus interval) from control (black) and PNE (red) in response to afferent fiber stimulation. **B.** The amplitude of the first EPSC is significantly reduced in PNE mice (Mann-Whitney, p=0.001, n=14 cells/group). **C.** Paired pulse ratio (PPR) is quantified from EPSC2/EPSC1. PPR is higher in PNE mice compared to control (Mann-Whitney, p=0.0002, n=14 cells/group). **D.** Representative traces of a 100 Hz train stimulation (500ms) from control (black) and PNE (red). **E.** EPSC amplitudes normalized to first EPSC during 100 Hz train, with significantly less depression observed in PNE mice compared to control (Two-way ANOVA, p=0.014, n=13-14 cells/group). **F.** Cumulative EPSC amplitudes are plotted across the 50 stimulations in the 100 Hz train. Y-intercepts (indicated by arrows) are estimates from linear regression backextrapolated using the steady state portion of the cumulative EPSC graph (last 10 points) to yield values proportional to RRP size. **G.** The Y-intercept is significantly reduced in PNE mice (Welch’s t-test, p=0.02, n=12-14 cells/group). The RRP, calculated from the Y-intercept/mEPSC amplitudes, shows no significant difference in vesicle number between groups (Welch’s t-test, p=0.15, n=12-14 cells/group). Probability of release (*P_r_*) is determined by dividing the first EPSC amplitude by the cumulative EPSC amplitude, revealing a significant decrease in *P_r_* in PNE mice compared to control (Welch’s t-test, p=0.028, n=11-13 cells/group).

Furthermore, we analyzed the readily releasable pool (RRP) size and probability of release (*P_r_*) using evoked EPSC trains at 100 Hz stimulation. During a 100 Hz stimulation train, the amplitudes of evoked EPSCs were gradually decreased in both control and PNE mice, referred to as short-term depression. When normalizing to the first EPSC amplitude, we found significantly less depression in PNE compared to control (**Figure 6D-E**, Two-way ANOVA, p=0.014, n=13-14 cells/group). To estimate RRP size and *P_r_*, we used the Schneggenburger-Meyer-Neher (SMN) method^27^, which plots cumulative EPSC amplitude against the stimulation number. A linear regression fitted to the steady state (last 10 EPSCs) is backextrapolated to the y-axis to give a y-intercept value which is proportional to RRP (**Figure 6F**). The Y-intercept values were significantly reduced in PNE mice (Control: 11.49 ± 1.64 nA vs. PNE: 6.58 ± 0.99 nA, Welch’s t-test, p=0.02, n=12-14 cells/group). To estimate readily releasable pool size, we divided the Y-intercept value by the average group mEPSC amplitude to yield an approximate number of glutamate vesicles per calyx terminal. After making this correction, we observed no significant difference in RRP between groups (**Figure 6G**, Control: 285.1 ± 40.79 vesicles vs PNE: 208.5 ± 31.36 vesicles, Welch’s t-test, p=0.15, n=12-14 cells/group). *P_r_* is calculated by dividing the Y-intercept by the first eEPSC amplitude. In PNE mice, *P_r_* of glutamate vesicles at the calyx terminal was significantly lower compared to control (PNE: 0.23 ± 0.01 vs Control: 0.29 ± 0.02, Welch’s t-test, p=0.028, n=11-13 cells/group). Overall, the results demonstrate that PNE impairs the development of glutamatergic synapses through alterations in both presynaptic and postsynaptic properties.

### Effects of PNE on auditory brainstem responses and cochlear function in mice

Next, we investigated the systemic impact of alterations in nAChR expression and synaptic transmission resulting from PNE on *in vivo* auditory function. To gain a comprehensive understanding of the relationship between PNE-induced alterations at the cellular level and the effects on auditory processing, we utilized the auditory brainstem response (ABR) test. In response to a sound stimulus, an experimenter will observe 5 distinct waves of neuronal activity corresponding to: I) auditory nerve, II) cochlear nucleus, III) superior olivary complex, IV) inferior colliculus, and V) medial geniculate nucleus. We assessed auditory function in both control and PNE animals at p21 by measuring ABRs from each group (**Figure 7A**, control in black, PNE in red). Specifically, we evaluated ABR thresholds, as well as the amplitudes and latencies of ABR waves, between the two groups. There was a significant increase in ABR click threshold in the PNE group compared to the control group (**Figure 7B**, Mann-Whitney test, p=0.021, n=11-14 mice/group). The elevated ABR threshold indicates PNE caused a reduction in the sensitivity of the auditory system to sound. To further understand the impact of PNE on auditory processing, we quantified the amplitudes and latencies of each peak in the ABR. Our analysis revealed that at 80 dB, the peak amplitudes of the ABR were significantly lower in PNE mice compared to control by two-way ANOVA (**Figure 7C**, p=0.029, n=11-14 mice/group). This result suggests a decrease in sound-evoked neuronal activity in each auditory nucleus in response to a click stimulus. However, there was no effect of treatment on ABR peak latency indicating no effect on central conduction speed (**Figure 7D**, Two-way ANOVA, p=0.07, n=11-1 animals/group). Moreover, we sought to determine whether the observed changes in ABRs were solely due to central processing deficits, or if the peripheral auditory system was also affected by PNE. To assess outer hair cell function in the cochlea, we used the distortion product otoacoustic emission (DPOAE) test. There was no significant difference in distortion product threshold between control and PNE mice (**Figure 7E**, Two-way ANOVA, p=0.60, n=12-14 mice/group) or distortion product amplitude (**Figure 7F**, Two-way ANOVA, p=0.39), indicating no significant impact of PNE on cochlea function. These results suggest that the observed ABR phenotypes were likely due to central processing deficits rather than peripheral auditory dysfunction. More specifically, PNE has significant impacts on the auditory brainstem rather than the peripheral system. Taken together, our results provide evidence that PNE can induce significant changes in auditory function at the cellular level, ultimately impacting auditory development.

**Figure 7.**
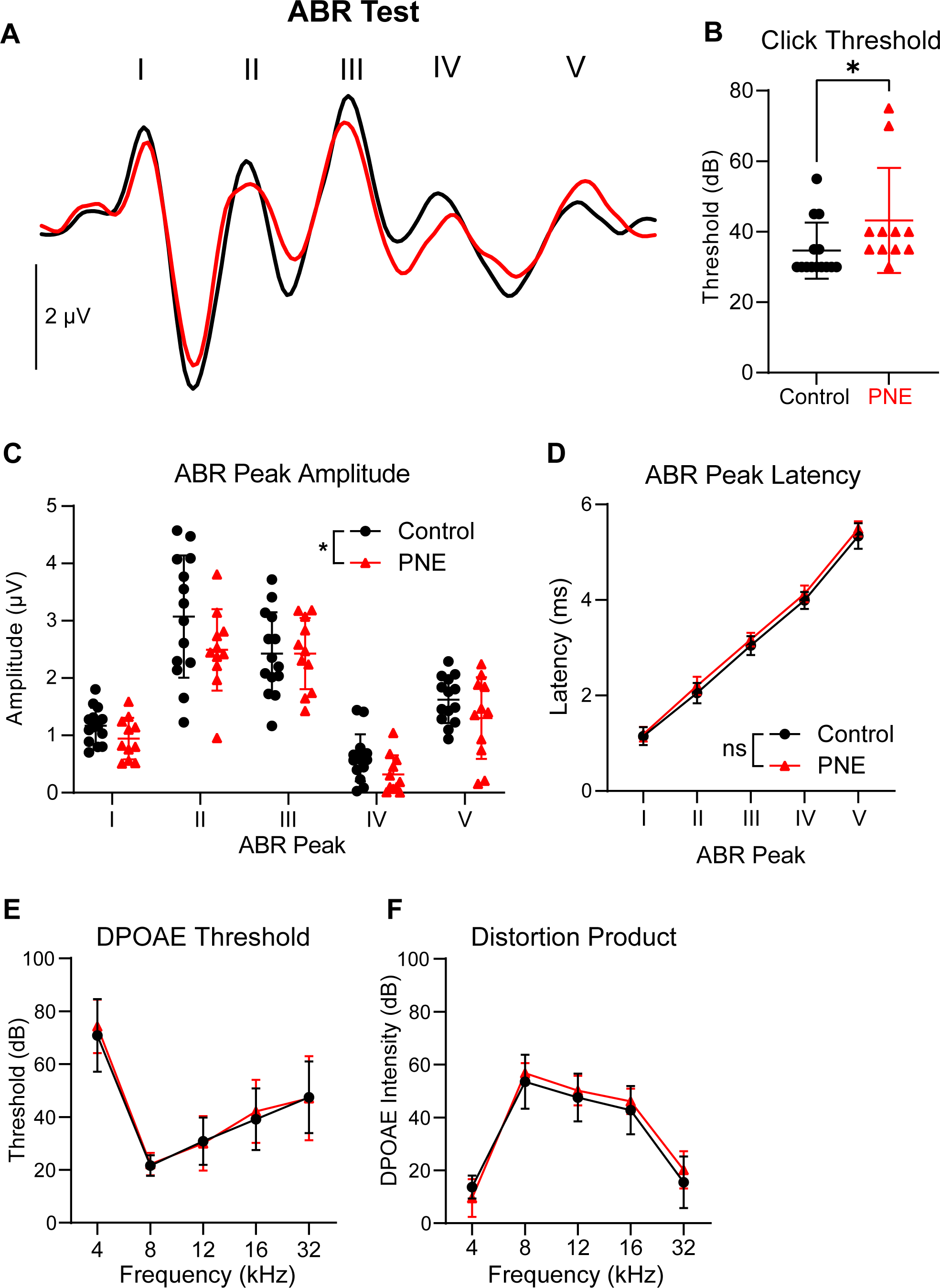
Impact of PNE on auditory brainstem responses and cochlear function in mice. **A.** Representative auditory brainstem response (ABR) traces evoked by click stimuli at 90 dB SPL from control (black) and PNE (red) mice at p21. **B.** A summary of ABR click thresholds in control and PNE mice, demonstrating a significant increase in ABR threshold in PNE mice (Mann Whitney test, p=0.021, n=11-14 mice/group). **C.** ABR amplitudes measured at 80 dB SPL in control and PNE mice, with PNE inducing a significant reduction in ABR amplitude compared to controls (Two-way ANOVA, p=0.029, n=11-14 mice/group). **D.** Peak latencies of ABR waves at 80 dB SPL in control and PNE mice, showing no significant effect of treatment on ABR wave peak latencies (Two-way ANOVA, p=0.07, n=11-14 mice/group). All bars in graphs represent mean ± S.D. **E.** Using DPOAE test, threshold sound intensity required to detect a distortion product was measured across frequencies. PNE did not significantly affect the threshold intensity required to detect a distortion product (DP, Two-way ANOVA, p=0.60, n=12-14 mice/group). **F.** DPs generated during DPOAE test, where PNE does not significantly impact the amplitude of DPs (Two-way ANOVA, p=0.39, n=12-14 mice/group).

## Discussion

Our study uncovers the adverse effects of PNE on the glutamatergic synaptic mechanisms and auditory processing in juvenile mice, highlighting the critical alterations in nAChRs function within the MNTB. We found that a diminished functional expression of postsynaptic nAChRs during the early postnatal period, while presynaptic nAChR activity, which is key for glutamate release, becomes more prominent after the onset of hearing. PNE disrupt this developmentally regulated pattern of nAChR expression, leading to impaired synaptic transmission at the calyx of Held-MNTB synapse. Additionally, PNE mice exhibit increased auditory brainstem response thresholds and reduced peak amplitudes, indicating impaired central auditory processing without cochlear function changes.

### Diversity of nAChR Expression and Function in the CNS

Previous studies indicate that nAChR subunit expression patterns vary across brain regions during postnatal development, with distinct developmental patterns observed in the brainstem, cerebellum, and striatum^28–31^. For example, α7 subunit mRNAs display consistent expression in the brainstem, but a bell-shaped developmental pattern in the striatum. In the auditory system, nAChR expression is initially widespread in the cochlear nucleus and superior olivary complex but becomes more restricted to specific cell types and synaptic compartments during development^9,15^. Our study revealed that nAChRs are expressed in both pre- and postsynaptic compartments of the MNTB synapse with discrete developmental patterning. It is possible that pre- and postsynaptic nAChRs have different expression timing due to differences in subunit composition^28,31^.

Functional differences in nAChR subunit composition allow for discrete pharmacological and physiological characteristics. Our data demonstrated that α4β2 nAChRs are responsible for a significant portion of nAChR currents from postsynaptic MNTB neurons. α4β2 nAChRs are the most prevalent nAChR subunit conformation in the brain which exhibit slow channel kinetics, low calcium permeability, and soma/postsynaptic density localization^32^. Due to their high affinity for nicotine, β2-containing nAChRs are required for mediating the reinforcing effects of nicotine^33,34^. The developmental role of α4β2 nAChRs is less well characterized, but maturation and patterning of retinothalamic projections require periodic waves of action potentials that are dependent on β2-containing nAChRs^35,36^. The role of α4β2 nAChRs in MNTB neuron development has yet to be explored. Due to their diverse functional profiles, MNTB principal neurons may express specific nAChR subtypes to stabilize calyx-MNTB synapse formation and subsequently downregulate or transition to another subunit composition once synapses are structurally mature around p14^37^.

In the adult brain, presynaptic nAChR localization is prevalent at glutamatergic synapses, such as terminals in the ventral tegmental area and hippocampus^38,39^. Although we cannot definitively conclude the role of presynaptic nAChRs in neurotransmitter release at the calyx terminal, we hypothesize they respond to top-down cholinergic inputs from the pontomesencephalic tegmentum (PMT) or collaterals from superior olivary complex (SOC) neurons to modulate MNTB activity^9^. Our data show that α7 nAChR activation regulates vesicular glutamate release from the calyx terminal in post-hearing animals. α7 nAChRs typically have fast channel kinetics, high calcium permeability, and presynaptic localization, making them well-suited for modulating vesicular release^40,41^. In the developing auditory cortex, α7 nAChR-dependent enhancement of NMDA-mediated neurotransmission contributes to experience-dependent refinement of synapses^11^. Taken together with our findings, cholinergic inputs to the MNTB enhance vesicular glutamate release from the calyx via α7 nAChRs to modulate synaptic transmission after hearing onset.

### Effect of PNE on nAChR patterning and glutamatergic neurotransmission

PNE has been shown to impair the maturation of glutamatergic neurotransmission in the brainstem in globular cells of the VNLL^21^. Our study found that PNE caused a significant decrease in functional presynaptic nAChRs in at the calyx terminal, but interestingly, a significant increase in postsynaptic nAChR-evoked currents in MNTB neurons at p20-21. An increase in functional postsynaptic nAChRs in the PNE is supported by the observed increased nAChR expression across human and animal models alike in response to chronic nicotine exposure^26,42–44^. Our results showed that PNE disrupted or delayed developmental changes in nAChR expression at pre- and postsynaptic compartments, which could have lasting effects on glutamate signaling and brain development. It remains to be determined whether this postsynaptic effect persists into adulthood or if the increase in nAChR-mediated currents is transient.

Impaired glutamatergic signaling following PNE is demonstrated by reduced mEPSC and eEPSC amplitudes but no change in RRP size, suggesting decreases in quantal size but no change in quantal content. (**Figure 6**). It is possible that the impaired calyx-MNTB neurotransmission is a secondary effect of disrupted nAChR patterning. PNE has been shown to alter the expression and function of nAChRs across brain regions, leading to changes in presynaptic release and glutamatergic neurotransmission^45–48^. Genetic deletion of α7 nAChRs decreases mEPSC frequency in the cortex and hippocampal neurons, indicating reduced number of glutamatergic synapses^49,50^. Additionally, PNE could impact presynaptic glutamate release through alterations in calcium signaling in the hippocampus and cortex^51–53^. Acute nAChR activation increases spontaneous glutamate vesicle fusion^47,54–58^, and specifically increases release probability demonstrated by decreased paired-pulse ratio (PPR) in hippocampal neurons^59–61^. Our study showed acute ACh application increase glutamate release, chronic nAChR activation in PNE mice reduced release probability (**Figure 3, 6**). Therefore, if PNE decreases presynaptic nAChR expression or its activity, this could contribute to decreased release probability of vesicular glutamate observed at the calyx terminal.

Moreover, PNE could modulate synaptic transmission by decreasing AMPAR expression in the postsynaptic membrane. Genetic deletion of neuroligin 3, a protein key for AMPAR stabilization in the postsynaptic membrane, has been shown to decrease eEPSC amplitude at the calyx terminal, similar to our PNE model^62^. PNE-induced decrease in mEPSC and eEPSC amplitude might be driven by both postsynaptic changes (e.g., a decrease in AMPAR expression, density, or clustering) and presynaptic mechanisms. Previous work observed a reduction in VGLUT1 and PSD95 in hippocampal synaptosomes following prenatal nicotine exposure^48^. PNE has been shown alter both presynaptic release mechanisms and postsynaptic receptor expression, leading to long-lasting changes in glutamate signaling and subsequent effects on brain development and function.

### Physiological Relevance

PNE-induced impairment of glutamatergic neurotransmission and altered nAChR expression in the MNTB might have important physiological consequences for auditory processing. The calyx of Held-MNTB synapse is critical for precise encoding of sound onset time, and intensity, and sound localization^63–65^. Our data reveal that PNE mice exhibit an elevated click ABR threshold and decreased ABR peak amplitudes, indicating a deficit in central auditory processing. The observed cellular phenotypes impact sound-evoked neuronal population activity via ABR, impairing crucial auditory functions. Additionally, alterations in presynaptic release probability and readily releasable pool size could have implications for short-term plasticity, which is thought to play a role in processing complex sounds such as speech^66,67^.

The disruptions in glutamatergic signaling and nAChR expression following PNE could have broader implications for central auditory function, given the widespread distribution of glutamatergic synapses in the auditory brainstem pathway^68^. Disruptions in nAChR signaling have been linked to impaired auditory processing in various conditions, such as schizophrenia-related auditory sensory gating deficits, which can be improved with nAChR agonists^69^. Similarly, genetic deletion of α9 nAChR in mice leads to impairment in sound localization and frequency discrimination^70^. On the other hand, artificially augmenting nAChR signaling to enhance medial olivocochlear efferent activity impairs glutamatergic neurotransmission at the calyx-MNTB synapse and eliminates tonotopic patterning of intrinsic properties of MNTB neurons^71^. Interestingly, our previous work has observed elimination of MNTB tonotopic gradients in mice with global BDNF reduction^72^, and neonatal exposure to nicotine reduces brain BDNF levels^73^. Therefore, it is possible that reduction of BDNF following chronic nicotine exposure is contributing to the PNE-induced synaptic impairments. In summary, our findings provide novel insights into the underlying mechanisms of auditory deficits following chronic nicotine exposure during early development and emphasize the importance of addressing maternal nicotine intake during pregnancy to mitigate the risk of auditory processing deficits in children.

## Materials and Methods

### Animals

Both sexes of C57BL/6J mice were used under the guidelines approved by the UT Health San Antonio Institutional Animal Care and Use Committee. All experiments were done between postnatal day 8 to 24 during the animals’ light cycle. Animals were housed in a 12-hr light/dark cycle.

### Perinatal Nicotine Exposure Protocol

Our perinatal nicotine exposure protocol utilizes subcutaneous injections of nicotine hydrogen tartrate (0.7 mg/kg free base nicotine) dissolved in 0.9% sterile saline to pups postnatal day 8 to 12 twice daily (totaling 1.4 mg/kg/day). Littermate controls were injected with comparable volumes of saline vehicle. Pups were taken from home cage for injections and monitored for 15 minutes post-injection before being returned to their home cage. Dosage has been used previously to study chronic nicotine effects on the brain, and on the auditory system specifically, to approximate blood levels of nicotine in smokers^7,74^.

### Ex Vivo Brain Slice Electrophysiology

#### Slice Preparation

Animals were anesthetized with isoflurane and quickly decapitated for brain slice collection. Brains were immediately removed and immersed in ice-cold low-calcium artificial cerebrospinal fluid (aCSF) containing (in mM): 125 NaCl, 2.5 KCl, 3 MgCl_2_, 0.1 CaCl_2_, 25 glucose, 25 NaHCO_3_, 1.25 NaH_2_PO_4_, at pH 7.4 and osmolarity 310–320 mOsm, bubbled with carbogen (95% O_2_, 5% CO_2_). Transverse 200 μm-thick brainstem slices containing the MNTB were collected using a Vibratome (VT1200S, Leica, Germany). After slices were collected, the slices were placed in an incubation chamber containing “reserve” aCSF bubbled with carbogen at 35°C for 30 min and then kept at room temperature for the remainder of the day. Reserve aCSF contains (in mM): 100 NaCl, 2.5 KCl, 2 MgCl_2_, 1 CaCl_2_, 25 glucose, 30 sucrose, 25 NaHCO_3_, 1.25 NaH_2_PO_4_ with similar pH and osmolarity to low calcium aCSF.

#### Electrophysiological Recordings

Whole-cell patch clamp recordings were done in room temperature normal aCSF, which is comprised of the same ingredients as low-calcium aCSF except it contains 1 mM MgCl_2_ and 2 mM CaCl_2_. Reagents utilized externally (in aCSF) and internally (pipette solution) for each experiment are described below. Figures 1-5 used K-gluconate-based internal which contains (in mM): 125 K-gluconate, 20 KCl, 5 Na_2_-phosphocreatine, 10 HEPES, 4 Mg-ATP, 0.2 EGTA, and 0.3 GTP, pH adjusted to 7.3 with KOH, osmolarity at 290 mOsm. Figure 6 used Cs-methane-sulfonate-based internal which contains (in mM): 130 Cs-methane-sulfonate, 10 CsCl, 5 EGTA, 10 HEPES, 4 ATP, 0.3 GTP, 5 Na_2_-phosphocreatine, 10 TEA-Cl, 2 QX 314 with osmolarity at 290 mOsm. Glass capillaries were pulled with Model P-1000 Micropipette Puller (Sutter Instruments) to obtain tip resistance at 3-6 MΩ. Voltage-clamp and current clamp-experiments were done with an EPC-10 amplifier (HEKA Electronik, Lambrecht, Pfalz, Germany). For postsynaptic voltage clamp experiments, holding potential was −65 mV. Series resistance was <20 MΩ without compensation, and if SR was >20MΩ, compensation up to 50% was utilized. Any cell with series resistance above 40MΩ was excluded from analysis. All mEPSC recordings had TTX (0.5 μM) and atropine (1 μM) externally, and calcium free mEPSC experiments had 0 mM CaCl_2_ and 4 mM MgCl_2_ externally. Acute puff application of ACh (1 mM) was performed with manual pressure using a 1 mL syringe for 1 second, and for Figure 2, using a Pneumatic Drug Ejection System from npi Electronic Instruments (10 psi for 500 ms). In Figure 2, postsynaptic pharmacology experiments utilized nAChR antagonists dihydro-β-erythroidine hydrobromide (DHβE, 3 μM, Tocris) and methyllycaconitine (MLA, 10 nM, Tocris). Drugs were bath-applied for 9 minutes before second puff application of ACh (4 mins for drug to reach bath then drug in contact with slice for 5 mins). When utilizing MLA for Figure 3, 10 nM MLA was in external aCSF for the entirety of recording experiments. For evoked EPSC recordings, electrical stimulation was generated with a bipolar stimulating electrode placed at midline. After determining threshold voltage of stimulator to elicit eEPSC, no more than 140% of threshold strength was used for the duration of the recording. Evoked EPSC recordings contained strychnine (2 μM) and bicuculline (20 μM) in the external solution.

#### Electrophysiological Data Analysis

Whole-cell recordings were collected with PatchMaster and exported to Igor Pro 8 (Wavemetrics, Lake Oswego, OR, United States) for analysis. For acute puff experiments, no more than 4 cells were recorded from per slice (alternating recordings with contralateral MNTB) to avoid desensitization of nAChRs. For mEPSC analysis, 30 seconds of activity pre- and post-ACh application was analyzed per cell with Mini Analysis (Synaptosoft). For evoked EPSCs, an average of 5 traces for single and paired pulse protocols was analyzed per cell. 100 Hz trains were averages of 2-3 traces for analysis. Probability of release (P_r_) and readily releasable pool (RRP) estimates were conducted with the Schneggenburger-Meyer-Neher Method^27^.

### *In vivo* Auditory Testing

Auditory testing took place in a sound-attenuated chamber (Med Associates, Albans, VT) with mice anesthetized with isoflurane (Vet One, India) levels controlled by a Vapomatic chamber (A.M. Bickford Inc., Wales Center, NY). Mice were initially anesthetized at 3.5% isoflurane and then maintained at 2.5% isoflurane (1 l/min O_2_ flow rate) during recordings. Body temperature was maintained around 37°C with an electric heating pad on the underside of the mouse. All acoustic stimuli were generated by an auditory evoked potentials workstation [Tucker-Davis Technologies (TDT), Alachua, FL, United States].

#### Auditory Brainstem Response (ABR) test

Subdermal needle electrodes (Rochester Electro-Medical, Lutz, FL, United States) were placed on the top of the head between the ears (active), ipsilateral mastoid (reference), and contralateral mastoid (ground) while animal was in prone position. Closed-field click stimulus was delivered to the left ear through a 10-cm plastic tube (Tygon; 3.2-mm outer diameter) from a Multi-Field Magnetic Speaker (Tucker Davis Technologies [TDT], Alachua, FL) at a repeat rate of 16/s. Sound intensities ranged from 90 to 20 dB, with 5-dB decrements. All analyses were done at 80 dB intensity. ABR wave amplitudes were analyzed from trough to peak.

#### Distortion product otoacoustic emission (DPOAE) test

DPOAE tests were performed similar to recent work from our lab^75^. Acoustic sound stimuli were delivered through two Multi-Field Magnetic Speakers in 10-cm coupling tubes (Tucker Davis Technologies [TDT], Alachua, FL) and connected with an ER-10B+ microphone with an ear tip (Etymotic Research, Elk Grove Village, IL) that were inserted into the mouse’s left ear canal. Pure tones were presented at 20% frequency separation between f1 and f2 at 4, 8, 12, 16 and 32 kHz, with intensities starting from 80 to 20 dB, decreasing by 10 dB at each presentation. 512 sweeps were averaged by the RZ6 processor (TDT), and distortion products were calculated as the noise floor subtracted from 2f1-f2.

### Statistical Analysis

All statistical analyses were performed in GraphPad Prism version 9.2.0 for Windows (GraphPad Software, San Diego, CA, United States). The normality of datasets was analyzed using the Kolmogorov-Smirnov test. Parametric or non-parametric tests were carried out accordingly. To compare two groups, an unpaired *t*-test with Welch’s correction (parametric) or Mann-Whitney U test (non-parametric) was carried out. For mEPSC analysis comparing frequency and amplitude before and after ACh application, paired t-tests or Wilcoxon paired-tests were used. To compare current amplitudes across ages, one-way or two-way ANOVAs were used. Figures 1 and 7 represent data as mean ± SD in figures and results. Figures 2-6 represent data as mean ± SEM in figures and results. Linear regressions done in Supplemental Figure 1 utilize t-tests to test if slopes are significantly non-zero.

## Acknowledgements

This work was supported by grants from the National Institute of Health: R01 DC018797(Kim), R01 NS123933 (Pugh), and F31DC021102 (Wollet).

---

**Figure S1.**
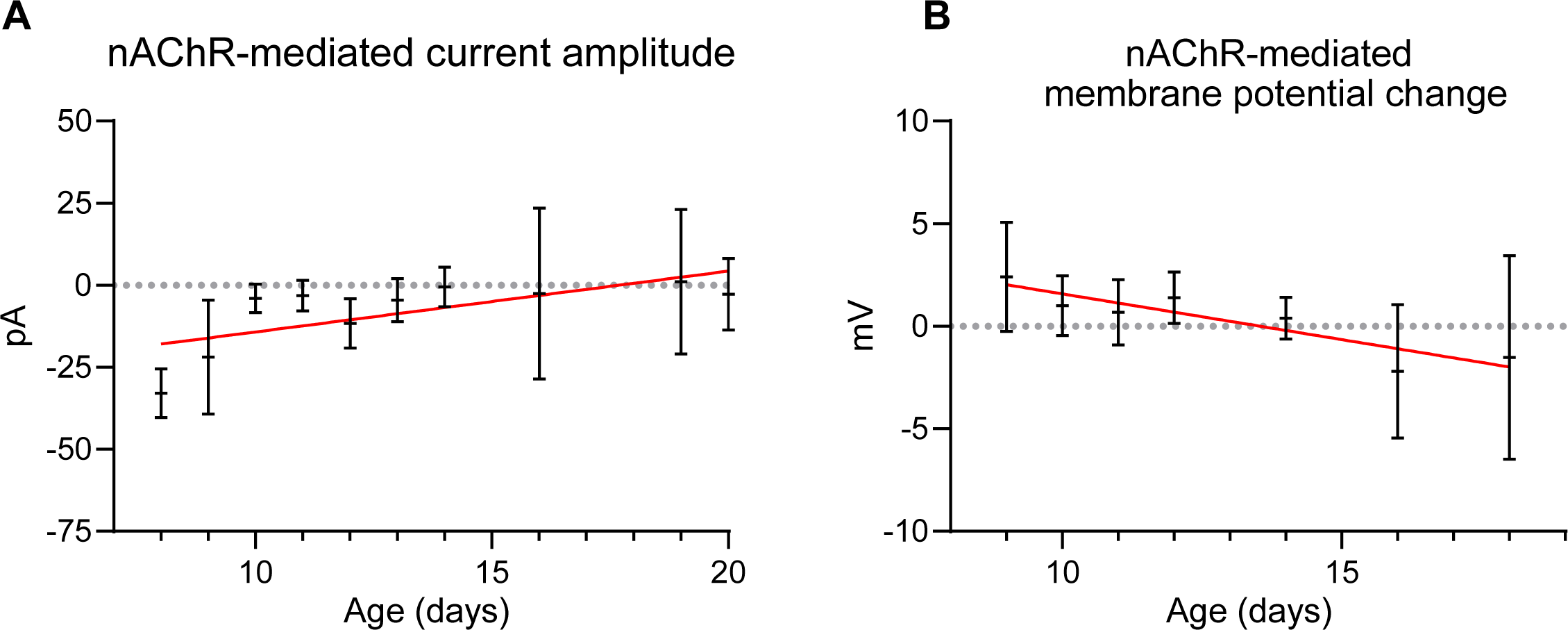
Developmental decline of functional nAChR expression in postnatal ages. **A.** nAChR-mediated currents plotted across postnatal age from Figure 1. Simple linear regression slope (red) is significantly non-zero (T-test, p=0.026). **B.** nAChR-mediated membrane potential changes plotted across postnatal age from Figure 1. Simple linear regression slope is significantly non-zero (T-test, p=0.005).

## References

1. King, E., Campbell, A., Belger, A., & Grewen, K. (2018). Prenatal Nicotine Exposure Disrupts Infant Neural Markers of Orienting. Nicotine & Tobacco Research, 20(7), 897–902. 10.1093/ntr/ntx177

2. Katbamna, B., Klutz, N., Pudrith, C., Lavery, J. P., & Ide, C. F. (2013). Prenatal smoke exposure: Effects on infant auditory system and placental gene expression. Neurotoxicology and Teratology, 38, 61–71. 10.1016/j.ntt.2013.04.008

3. Crane, L., Goddard, L., & Pring, L. (2009). Sensory processing in adults with autism spectrum disorders. Autism, 13(3), 215–228. 10.1177/1362361309103794

4. Marco, E. J., Hinkley, L. B. N., Hill, S. S., & Nagarajan, S. S. (2011). Sensory Processing in Autism: A Review of Neurophysiologic Findings: *Pediatric Research*, 69(5 Part 2), 48R-54R. 10.1203/PDR.0b013e3182130c54

5. Ghanizadeh, A. (2011). Sensory Processing Problems in Children with ADHD, a Systematic Review. Psychiatry Investigation, 8(2), 89. 10.4306/pi.2011.8.2.89

6. Kramarow, E., & Elgaddal, N. (2023). Current Electronic Cigarette Use in Adults Aged 18 and Over: United States, 2021. National Center for Health Statistics (U.S.). 10.15620/cdc:129966

7. Aramakis, V. B., Hsieh, C. Y., Leslie, F. M., & Metherate, R. (2000). A Critical Period for Nicotine-Induced Disruption of Synaptic Development in Rat Auditory Cortex. The Journal of Neuroscience, 20(16), 6106–6116. 10.1523/JNEUROSCI.20-16-06106.2000

8. Sun, W., Hansen, A., Zhang, L., Lu, J., Stolzberg, D., & Kraus, K. S. (2008). Neonatal nicotine exposure impairs development of auditory temporal processing. Hearing Research, 245(1–2), 58–64. 10.1016/j.heares.2008.08.009

9. Zhang, C., Beebe, N. L., Schofield, B. R., Pecka, M., & Burger, R. M. (2021). Endogenous Cholinergic Signaling Modulates Sound-Evoked Responses of the Medial Nucleus of the Trapezoid Body. The Journal of Neuroscience, 41(4), 674– 688. 10.1523/JNEUROSCI.1633-20.2020

10. Beebe, N. L., Zhang, C., Burger, R. M., & Schofield, B. R. (2021). Multiple Sources of Cholinergic Input to the Superior Olivary Complex. Frontiers in Neural Circuits, 15, 715369. 10.3389/fncir.2021.715369

11. Aramakis, V. B., & Metherate, R. (1998). Nicotine Selectively Enhances NMDA Receptor-Mediated Synaptic Transmission during Postnatal Development in Sensory Neocortex. The Journal of Neuroscience, 18(20), 8485–8495. 10.1523/JNEUROSCI.18-20-08485.1998

12. Elgoyhen, A. B., Katz, E., & Fuchs, P. A. (2009). The nicotinic receptor of cochlear hair cells: A possible pharmacotherapeutic target? Biochemical Pharmacology, 78(7), 712–719. 10.1016/j.bcp.2009.05.023

13. Happe, H. K., & Morley, B. J. (1998). Nicotinic acetylcholine receptors in rat cochlear nucleus: [125I]-?-bungarotoxin receptor autoradiography and in situ hybridization ofL?7 nAChR subunit mRNA. The Journal of Comparative Neurology, 397(2), 163–180. 10.1002/(SICI)1096-9861(19980727)397:2<163::AID-CNE2>3.0.CO;2-Z

14. Yao, W., & Godfrey, D. A. (1999). Immunolocalization of α4 and α7 subunits of nicotinic receptor in rat cochlear nucleus. Hearing Research, 128(1–2), 97–102. 10.1016/S0378-5955(98)00199-3

15. Happe, H. K., & Morley, B. J. (2004). Distribution and postnatal development of α7 nicotinic acetylcholine receptors in the rodent lower auditory brainstem. Developmental Brain Research, 153(1), 29–37. 10.1016/j.devbrainres.2004.07.004

16. Ghimire, M., Cai, R., Ling, L., Hackett, T. A., & Caspary, D. M. (2020). Nicotinic Receptor Subunit Distribution in Auditory Cortex: Impact of Aging on Receptor Number and Function. The Journal of Neuroscience, 40(30), 5724–5739. 10.1523/JNEUROSCI.0093-20.2020

17. Metherate, R., Intskirveli, I., & Kawai, H. D. (2012). Nicotinic filtering of sensory processing in auditory cortex. Frontiers in Behavioral Neuroscience, 6. 10.3389/fnbeh.2012.00044

18. Chen, G., & Yan, J. (2007). Cholinergic modulation incorporated with a tone presentation induces frequency-specific threshold decreases in the auditory cortex of the mouse: Cholinergic facilitation for cortical plasticity. European Journal of Neuroscience, 25(6), 1793–1803. 10.1111/j.1460-9568.2007.05432.x

19. Rivera-Perez, L. M., Kwapiszewski, J. T., & Roberts, M. T. (2021). Α3β4∗ Nicotinic Acetylcholine Receptors Strongly Modulate the Excitability of VIP Neurons in the Mouse Inferior Colliculus. Frontiers in Neural Circuits, 15, 709387. 10.3389/fncir.2021.709387

20. Weimann, S. R., Zhang, C., & Burger, R. M. (2023). A Developmental Switch in Cholinergic Mechanisms of Modulation in the Medial Nucleus of the Trapezoid Body. *The Journal of Neuroscience*, JN-RM-0356–23. 10.1523/JNEUROSCI.0356-23.2023

21. Baumann, V. J., & Koch, U. (2017). Perinatal nicotine exposure impairs the maturation of glutamatergic inputs in the auditory brainstem: Nicotine exposure disturbs maturation of synaptic inputs. The Journal of Physiology, 595(11), 3573– 3590. 10.1113/JP274059

22. Giniatullin, R., Nistri, A., & Yakel, J. (2005). Desensitization of nicotinic ACh receptors: Shaping cholinergic signaling. Trends in Neurosciences, 28(7), 371–378. 10.1016/j.tins.2005.04.009

23. Paradiso, K. G., & Steinbach, J. H. (2003). Nicotine is Highly Effective at Producing Desensitization of Rat α4β2 Neuronal Nicotinic Receptors. The Journal of Physiology, 553(3), 857–871. 10.1113/jphysiol.2003.053447

24. Fenster, C. P., Whitworth, T. L., Sheffield, E. B., Quick, M. W., & Lester, R. A. J. (1999). Upregulation of Surface α4β2 Nicotinic Receptors Is Initiated by Receptor Desensitization after Chronic Exposure to Nicotine. The Journal of Neuroscience, 19(12), 4804–4814. 10.1523/JNEUROSCI.19-12-04804.1999

25. Hsieh, C. Y., Leslie, F. M., & Metherate, R. (2002). Nicotine exposure during a postnatal critical period alters NR2A and NR2B mRNA expression in rat auditory forebrain. Developmental Brain Research, 133(1), 19–25. 10.1016/S0165-3806(01)00314-5

26. Vallejo, Y. F. (2005). Chronic Nicotine Exposure Upregulates Nicotinic Receptors by a Novel Mechanism. Journal of Neuroscience, 25(23), 5563–5572. 10.1523/JNEUROSCI.5240-04.2005

27. Schneggenburger, R., Meyer, A. C., & Neher, E. (1999). Released Fraction and Total Size of a Pool of Immediately Available Transmitter Quanta at a Calyx Synapse. Neuron, 23(2), 399–409. 10.1016/S0896-6273(00)80789-8

28. Zhang, X., Liu, C., Miao, H., Gong, Z., & Nordberg, A. (1998). Postnatal changes of nicotinic acetylcholine receptor α2, α3, α4, α7 and β2 subunits genes expression in rat brain. International Journal of Developmental Neuroscience, 16(6), 507–518. 10.1016/S0736-5748(98)00044-6

29. Son, J.-H., & Winzer-Serhan, U. H. (2006). Postnatal expression of α2 nicotinic acetylcholine receptor subunit mRNA in developing cortex and hippocampus. Journal of Chemical Neuroanatomy, 32(2–4), 179–190. 10.1016/j.jchemneu.2006.09.001

30. Shacka, J. J., & Robinson, S. E. (1998). Postnatal developmental regulation of neuronal nicotinic receptor subunit α7 and multiple α4 and β2 mRNA species in the rat. Developmental Brain Research, 109(1), 67–75. 10.1016/S0165-3806(98)00058-3

31. Azam, L., Chen, Y., & Leslie, F. M. (2007). Developmental regulation of nicotinic acetylcholine receptors within midbrain dopamine neurons. Neuroscience, 144(4), 1347–1360. 10.1016/j.neuroscience.2006.11.011

32. Posadas, I., Lopez-Hernandez, B., & Cena, V. (2013). Nicotinic Receptors in Neurodegeneration. Current Neuropharmacology, 11(3), 298–314. 10.2174/1570159X11311030005

33. Epping-Jordan, M. P., Picciotto, M. R., Changeux, J.-P., & Pich, E. M. (1999). Assessment of nicotinic acetylcholine receptor subunit contributions to nicotine self-administration in mutant mice. Psychopharmacology, 147(1), 25–26. 10.1007/s002130051135

34. Picciotto, M. R., Zoli, M., Rimondini, R., Léna, C., Marubio, L. M., Pich, E. M., Fuxe, K., & Changeux, J.-P. (1998). Acetylcholine receptors containing the β2 subunit are involved in the reinforcing properties of nicotine. Nature, 391(6663), 173–177. 10.1038/34413

35. Penn, A. A., Riquelme, P. A., Feller, M. B., & Shatz, C. J. (1998). Competition in Retinogeniculate Patterning Driven by Spontaneous Activity. Science, 279(5359), 2108–2112. 10.1126/science.279.5359.2108

36. Feller, M. B., Wellis, D. P., Stellwagen, D., Werblin, F. S., & Shatz, C. J. (1996). Requirement for Cholinergic Synaptic Transmission in the Propagation of Spontaneous Retinal Waves. Science, 272(5265), 1182–1187. 10.1126/science.272.5265.1182

37. Kandler, K., & Friauf, E. (1993). Pre- and postnatal development of efferent connections of the cochlear nucleus in the rat. The Journal of Comparative Neurology, 328(2), 161–184. 10.1002/cne.903280202

38. Jones, I. W., & Wonnacott, S. (2004). Precise Localization of α7 Nicotinic Acetylcholine Receptors on Glutamatergic Axon Terminals in the Rat Ventral Tegmental Area. The Journal of Neuroscience, 24(50), 11244–11252. 10.1523/JNEUROSCI.3009-04.2004

39. Fabian-Fine, R., Skehel, P., Errington, M. L., Davies, H. A., Sher, E., Stewart, M. G., & Fine, A. (2001). Ultrastructural Distribution of the α7 Nicotinic Acetylcholine Receptor Subunit in Rat Hippocampus. The Journal of Neuroscience, 21(20), 7993–8003. 10.1523/JNEUROSCI.21-20-07993.2001

40. Mansvelder, H. D., & McGehee, D. S. (2002). Cellular and synaptic mechanisms of nicotine addiction. Journal of Neurobiology, 53(4), 606–617. 10.1002/neu.10148

41. Dani, J. A. (2015). Neuronal Nicotinic Acetylcholine Receptor Structure and Function and Response to Nicotine. In International Review of Neurobiology (Vol. 124, pp. 3–19). Elsevier. 10.1016/bs.irn.2015.07.001

42. Breese, C. R., Marks, M. J., Logel, J., Adams, C. E., Sullivan, B., Collins, A. C., & Leonard, S. (1997). Effect of smoking history on [3H]nicotine binding in human postmortem brain. The Journal of Pharmacology and Experimental Therapeutics, 282(1), 7–13.

43. Perry, D. C., Dávila-García, M. I., Stockmeier, C. A., & Kellar, K. J. (1999). Increased nicotinic receptors in brains from smokers: Membrane binding and autoradiography studies. The Journal of Pharmacology and Experimental Therapeutics, 289(3), 1545–1552.

44. Harkness, P. C., & Millar, N. S. (2002). Changes in Conformation and Subcellular Distribution of α4β2 Nicotinic Acetylcholine Receptors Revealed by Chronic Nicotine Treatment and Expression of Subunit Chimeras. The Journal of Neuroscience, 22(23), 10172–10181. 10.1523/JNEUROSCI.22-23-10172.2002

45. Abdulla, F. A., Gray, J. A., Sinden, J. D., Bradbury, E., Calaminici, M.-R., Lippiello, P. M., & Wonnacott, S. (1996). Relationship between up-regulation of nicotine binding sites in rat brain and delayed cognitive enhancement observed after chronic or acute nicotinic receptor stimulation. Psychopharmacology, 124(4), 323–331. 10.1007/BF02247437

46. Pilarski, J. Q., Wakefield, H. E., Fuglevand, A. J., Levine, R. B., & Fregosi, R. F. (2012). Increased nicotinic receptor desensitization in hypoglossal motor neurons following chronic developmental nicotine exposure. Journal of Neurophysiology, 107(1), 257–264. 10.1152/jn.00623.2011

47. Proctor, W. R., Dobelis, P., Moritz, A. T., & Wu, P. H. (2011). Chronic nicotine treatment differentially modifies acute nicotine and alcohol actions on GABAA and glutamate receptors in hippocampal brain slices: Synaptic interactions between nicotine and alcohol. British Journal of Pharmacology, 162(6), 1351– 1363. 10.1111/j.1476-5381.2010.01141.x

48. Parameshwaran, K., Buabeid, M. A., Karuppagounder, S. S., Uthayathas, S., Thiruchelvam, K., Shonesy, B., Dityatev, A., Escobar, M. C., Dhanasekaran, M., & Suppiramaniam, V. (2012). Developmental nicotine exposure induced alterations in behavior and glutamate receptor function in hippocampus. Cellular and Molecular Life Sciences, 69(5), 829–841. 10.1007/s00018-011-0805-4

49. Lozada, A. F., Wang, X., Gounko, N. V., Massey, K. A., Duan, J., Liu, Z., & Berg, D. K. (2012). Glutamatergic Synapse Formation is Promoted by 7-Containing Nicotinic Acetylcholine Receptors. Journal of Neuroscience, 32(22), 7651–7661. 10.1523/JNEUROSCI.6246-11.2012

50. Zhong, C., Akmentin, W., Role, L. W., & Talmage, D. A. (2022). Axonal α7* nicotinic acetylcholine receptors modulate glutamatergic signaling and synaptic vesicle organization in ventral hippocampal projections. Frontiers in Neural Circuits, 16, 978837. 10.3389/fncir.2022.978837

51. Hayashida, S., Katsura, M., Torigoe, F., Tsujimura, A., & Ohkuma, S. (2005). Increased expression of L-type high voltage-gated calcium channel α1 and α2/δ subunits in mouse brain after chronic nicotine administration. Molecular Brain Research, 135(1–2), 280–284. 10.1016/j.molbrainres.2004.11.002

52. Katsura, M., Mohri, Y., Shuto, K., Hai-Du, Y., Amano, T., Tsujimura, A., Sasa, M., & Ohkuma, S. (2002). Up-regulation of L-type Voltage-dependent Calcium Channels after Long Term Exposure to Nicotine in Cerebral Cortical Neurons. Journal of Biological Chemistry, 277(10), 7979–7988. 10.1074/jbc.M109466200

53. Katsura, M., & Ohkuma, S. (2005). Functional Proteins Involved in Regulation of Intracellular Ca2+ for Drug Development: Chronic Nicotine Treatment Upregulates L-Type High Voltage-Gated Calcium Channels. Journal of Pharmacological Sciences, 97(3), 344–347. 10.1254/jphs.FMJ04007X3

54. Tang, B., Luo, D., Yang, J., Xu, X.-Y., Zhu, B.-L., Wang, X.-F., Yan, Z., & Chen, G.-J. (2015). Modulation of AMPA receptor mediated current by nicotinic acetylcholine receptor in layer I neurons of rat prefrontal cortex. Scientific Reports, 5(1), 14099. 10.1038/srep14099

55. 55. Le Magueresse, C., Safiulina, V., Changeux, J.-P., & Cherubini, E. (2006). Nicotinic modulation of network and synaptic transmission in the immature hippocampus investigated with genetically modified mice: Nicotine and synaptic transmission in the developing hippocampus. The Journal of Physiology, 576(2), 533–546. 10.1113/jphysiol.2006.117572

56. Sharma, G., & Vijayaraghavan, S. (2003). Modulation of Presynaptic Store Calcium Induces Release of Glutamate and Postsynaptic Firing. Neuron, 38(6), 929–939. 10.1016/S0896-6273(03)00322-2

57. Zhong, C., Du, C., Hancock, M., Mertz, M., Talmage, D. A., & Role, L. W. (2008). Presynaptic Type III Neuregulin 1 Is Required for Sustained Enhancement of Hippocampal Transmission by Nicotine and for Axonal Targeting of α7 Nicotinic Acetylcholine Receptors. The Journal of Neuroscience, 28(37), 9111–9116. 10.1523/JNEUROSCI.0381-08.2008

58. Puddifoot, C. A., Wu, M., Sung, R.-J., & Joiner, W. J. (2015). Ly6h Regulates Trafficking of Alpha7 Nicotinic Acetylcholine Receptors and Nicotine-Induced Potentiation of Glutamatergic Signaling. The Journal of Neuroscience, 35(8), 3420–3430. 10.1523/JNEUROSCI.3630-14.2015

59. Sola, E., Capsoni, S., Rosato-Siri, M., Cattaneo, A., & Cherubini, E. (2006). Failure of nicotine-dependent enhancement of synaptic efficacy at Schaffer-collateral CA1 synapses of AD11 anti-nerve growth factor transgenic mice. European Journal of Neuroscience, 24(5), 1252–1264. 10.1111/j.1460-9568.2006.04996.x

60. Cheng, Q., & Yakel, J. L. (2014). Presynaptic α7 Nicotinic Acetylcholine Receptors Enhance Hippocampal Mossy Fiber Glutamatergic Transmission via PKA Activation. The Journal of Neuroscience, 34(1), 124–133. 10.1523/JNEUROSCI.2973-13.2014

61. Damborsky, J. C., Griffith, W. H., & Winzer-Serhan, U. H. (2015). Neonatal nicotine exposure increases excitatory synaptic transmission and attenuates nicotine-stimulated GABA release in the adult rat hippocampus. Neuropharmacology, 88, 187–198. 10.1016/j.neuropharm.2014.06.010

62. Han, Y., Cao, R., Qin, L., Chen, L. Y., Tang, A.-H., Südhof, T. C., & Zhang, B. (2022). Neuroligin-3 confines AMPA receptors into nanoclusters, thereby controlling synaptic strength at the calyx of Held synapses. Science Advances, 8(24), eabo4173. 10.1126/sciadv.abo4173

63. Englitz, B., Tolnai, S., Typlt, M., Jost, J., & Rübsamen, R. (2009). Reliability of Synaptic Transmission at the Synapses of Held In Vivo under Acoustic Stimulation. PLoS ONE, 4(10), e7014. 10.1371/journal.pone.0007014

64. Guinan, J. J., & Li, R. Y.-S. (1990). Signal processing in brainstem auditory neurons which receive giant endings (calyces of Held) in the medial nucleus of the trapezoid body of the cat. Hearing Research, 49(1–3), 321–334. 10.1016/0378-5955(90)90111-2

65. Beiderbeck, B., Myoga, M. H., Müller, N. I. C., Callan, A. R., Friauf, E., Grothe, B., & Pecka, M. (2018). Precisely timed inhibition facilitates action potential firing for spatial coding in the auditory brainstem. Nature Communications, 9(1), 1771. 10.1038/s41467-018-04210-y

66. Singh, M., Miura, P., & Renden, R. (2018). Age-related defects in short-term plasticity are reversed by acetyl-L-carnitine at the mouse calyx of Held. Neurobiology of Aging, 67, 108–119. 10.1016/j.neurobiolaging.2018.03.015

67. Keine, C., Al-Yaari, M., Radulovic, T., Thomas, C. I., Valino Ramos, P., Guerrero-Given, D., Ranjan, M., Taschenberger, H., Kamasawa, N., & Young, S. M. (2022). Presynaptic Rac1 controls synaptic strength through the regulation of synaptic vesicle priming. eLife, 11, e81505. 10.7554/eLife.81505

68. Petralia, R. S., & Wenthold, R. J. (2009). Neurotransmitters in the Auditory System. In M. D. Binder, N. Hirokawa, & U. Windhorst (Eds.), Encyclopedia of Neuroscience (pp. 2847–2853). Springer Berlin Heidelberg. 10.1007/978-3-540-29678-2_3957

69. Aidelbaum, R., Labelle, A., Baddeley, A., & Knott, V. (2018). Assessing the acute effects of CDP-choline on sensory gating in schizophrenia: A pilot study. Journal of Psychopharmacology, 32(5), 541–551. 10.1177/0269881117746903

70. Clause, A., Lauer, A. M., & Kandler, K. (2017). Mice Lacking the Alpha9 Subunit of the Nicotinic Acetylcholine Receptor Exhibit Deficits in Frequency Difference Limens and Sound Localization. Frontiers in Cellular Neuroscience, 11, 167. 10.3389/fncel.2017.00167

71. 71. Di Guilmi, M. N., Boero, L. E., Castagna, V. C., Rodríguez-Contreras, A., Wedemeyer, C., Gómez-Casati, M. E., & Elgoyhen, A. B. (2019). Strengthening of the Efferent Olivocochlear System Leads to Synaptic Dysfunction and Tonotopy Disruption of a Central Auditory Nucleus. The Journal of Neuroscience, 39(36), 7037–7048. 10.1523/JNEUROSCI.2536-18.2019

72. Wollet, M., & Kim, J. H. (2022). Brain-Derived Neurotrophic Factor Is Involved in Activity-Dependent Tonotopic Refinement of MNTB Neurons. Frontiers in Neural Circuits, 16. https://www.frontiersin.org/article/10.3389/fncir.2022.784396

73. Xiaoyu, W. (2015). The exposure to nicotine affects expression of brain-derived neurotrophic factor (BDNF) and nerve growth factor (NGF) in neonate rats. Neurological Sciences, 36(2), 289–295. 10.1007/s10072-014-934-y

74. Liang, K., Poytress, B. S., Chen, Y., Leslie, F. M., Weinberger, N. M., & Metherate, R. (2006). Neonatal nicotine exposure impairs nicotinic enhancement of central auditory processing and auditory learning in adult rats. European Journal of Neuroscience, 24(3), 857–866. 10.1111/j.1460-9568.2006.04945.x

75. Nip, K., Kashiwagura, S., & Kim, J. H. (2022). Loss of β4-spectrin impairs Nav channel clustering at the heminode and temporal fidelity of presynaptic spikes in developing auditory brain. Scientific Reports, 12(1), 5854. 10.1038/s41598-022-09856-9

